# Assignment of molecular origins of NOE signal at −3.5 ppm in the brain

**DOI:** 10.1101/2023.02.03.526979

**Authors:** Yu Zhao, Casey Sun, Zhongliang Zu

## Abstract

**Purpose:** Nuclear Overhauser Enhancement mediated saturation transfer effect, termed NOE(−3.5 ppm), is a major source of chemical exchange saturation transfer (CEST) MRI contrasts at 3.5 ppm in the brain. Previous phantom experiments have demonstrated that both proteins and lipids, two major components in tissues, have substantial contributions to NOE(−3.5 ppm) signals. Their relative contributions in tissues are informative for the interpretation of NOE(−3.5 ppm) contrasts that could provide potential imaging biomarkers for relevant diseases, which remain incompletely understood.

**Methods:** Experiments on homogenates and supernatants of brain tissues collected from healthy rats, that could isolate proteins from lipids, were performed to evaluate the relative contribution of lipids to NOE(−3.5 ppm) signals. On the other hand, experiments on ghost membranes with varied pH, and reconstituted phospholipids with different chemical compositions were conducted to study the dependence of NOE(−3.5 ppm) on physiological conditions. Besides, CEST imaging on rat brains bearing 9L tumors and healthy rat brains was performed to analyze the causes of the NOE(−3.5 ppm) contrast variations between tumors and normal tissues, and between gray matter and white matter.

**Results:** Our experiments reveal that lipids have dominant contributions to the NOE (−3.5 ppm) signals. Further analysis suggests that decreased NOE(−3.5 ppm) signals in tumors and higher NOE(−3.5 ppm) signals in white matter than in gray matter are mainly explained by changes in membrane lipids, rather than proteins.

**Conclusion:** NOE(−3.5 ppm) could be exploited as a highly sensitive MRI contrast for imaging membrane lipids in the brain.

## INTRODUCTION

Chemical exchange saturation transfer (CEST) is an MRI contrast mechanism for indirectly detecting dilute solute molecules with enhanced sensitivity by measuring changes in water signals (1–5). In CEST imaging, a Z-spectrum is usually created by plotting water signals as a function of the frequency offset of RF saturation pulse, in which CEST effects sourced from multiple molecular pools can be investigated. For instance, in the central nervous system, amide proton transfer (APT) effect at 3.5 ppm, amine-water exchange effect at 3 ppm, guanidinium proton transfer effect at 2 ppm, Nuclear Overhauser Enhancement (NOE) medicated CEST effect at −1.6 ppm and −3.5 ppm (abbreviated as NOE(−1.6 ppm) and NOE(−3.5 ppm) in this study), along with a broad semi-solid MT effect can be observed in the Z-spectrum. Most of the CEST effects observed in the Z-spectrum have been assigned to certain molecular sources. Specifically, the APT effect originates from backbone amide protons on mobile proteins/peptides (6); the amine-water exchange effect originates from lysine protons on proteins, glutamate and free amino acids (7,8); the guanidinium proton transfer effect originates from creatine and arginine residues of mobile proteins (9–11); the NOE(−1.6 ppm) effect originates from choline head groups on phospholipids (12,13); the MT effect originates from immobile macromolecules (e.g. myelin) (14,15). In recent studies, although the origins of the NOE(−3.5 ppm) effect have been assigned to aliphatic protons associated with mobile proteins and lipids (12,13,16–19), their relative contributions to the *in vivo* NOE(−3.5 ppm) signals remain incompletely understood. Further studies on this issue would be highly rewarding since the NOE(−3.5 ppm) imaging has shown great potential for applications on brain tumors (12,17,20–24), Alzheimer’s disease (25) and spinal cord injury (26), etc.

NOE is a dipolar cross-relaxation between two nuclear spins at a close distance on a molecule (27–29). The cross-relaxation rate (or NOE coupling rate) depends on the speed of the molecule tumbling due to thermal motions. NOE usually are seen from macromolecules or molecules that have aliphatic protons with limited mobility. The proteins and lipids that are the major sources of the NOE signals play important roles in physiological functions of the brain. Proteins perform many functions in cells including catalyzing metabolic reactions and DNA replications, responding to stimuli, as well as providing structure to cells and organisms; lipids (i.e., phospholipids) are major components of cellular membranes, which are critical to maintaining the structure and functions of cells. Changes in the two molecules in specific diseases represent a different pathological process. Thus, further evaluation of their relative contributions to the NOE(−3.5 ppm) signal in biological tissues is informative for the interpretation of the NOE(−3.5 ppm) contrast.

Previously, the NOE(−3.5 ppm) signal from bovine serum albumin (BSA) phantoms (22,30) and reconstituted phospholipid phantoms (12,13) have been reported. Also, it was reported that NOE(−3.5 ppm) signal from proteins is sensitive to the structural integrity of proteins (22,30), and the NOE(−3.5 ppm) signal from phospholipids is sensitive to the spatial conformations and chemical compositions (12,13). However, the relative contributions of proteins and lipids to NOE(−3.5 ppm) signals in tissues cannot be assessed by these simple phantom experiments for the following reasons. First, BSA cannot represent the proteins in biological tissues that have different molecular structures and weights which could alter the NOE signals in human cells. Second, reconstituted phospholipids in the phantom experiments have a limited biophysical reality with respect to the cell membranes in tissues.

In this study, tissue homogenates are used to investigate the NOE(−3.5 ppm) effect to ensure a biophysical reality. Furthermore, the supernatant of the homogenates with the removal of most lipids is used to estimate the net contribution of the NOE(−3.5 ppm) effect from proteins. Although absolute concentrations of proteins in the supernatant may be lower than that in the tissue homogenates due to incomplete wall-broken of cells, the relative amplitudes of protein-sourced NOE(−3.5 ppm) signal and protein-sourced APT signal could keep unchanged since the cells with incomplete wall-broken have the similar protein components as that with complete wall-broken. Thus, a comparison of the ratio of the NOE(−3.5 ppm) signal to the APT signal between the tissue homogenates and the supernatant is performed to assess the relative contributions of proteins and lipids to the NOE(−3.5 ppm) signals. Besides, experiments on ghost membranes with varied pH, and reconstituted phospholipids with different chemical compositions were conducted to study the dependence of NOE(−3.5 ppm) on physiological conditions.

## MATERIALS AND METHODS

### Animal preparation

Three rats bearing 9L tumors and three healthy rats were included in this study. For brain tumor induction, each rat was injected with 1 × 10^5^ 9L glioblastoma cells in the right brain hemisphere and was then imaged after 2 to 3 weeks. During data acquisition, all rats were immobilized and anesthetized with a 2%/98% isoflurane/oxygen mixture. Respiration was monitored to be stable, and a constant rectal temperature of 37°C was maintained throughout the MRI imaging using a warm-air feedback system (SA Instruments, Stony Brook, NY, USA). All animal procedures were approved by the Animal Care and Usage Committee of Vanderbilt University Medical Center.

### Phantom preparation

The homogenates and the supernatant of brain tissues were prepared from two freshly sacrificed healthy rats. Brain tissues were weighed and washed quickly with 1 × phosphate buffered saline (PBS) to remove residual blood after being removed from the rats. Then, after adding 4-times 1 × PBS (w/w), the tissues were homogenized with a motor-driven blade-type homogenizer (Brinkmann Polytron PT3000; Kinematica, Malters, Switzerland) at a speed of 20,000 rpm for 2 × 30 second bursts. The homogenates were divided into two aliquots, one of which is used for the homogenate sample. The supernatant of brain tissues was separated from the other aliquot by removing sediments after centrifugation (20,000 × g, 30 min, 4°C). Finally, the two aliquots were titrated to 7.0 at the room temperature by using NaOH/HCl.

Ghost membranes were used to further investigate the origin of the NOE(−3.5 ppm) signal from the plasma membrane and to study the dependence of the NOE(−3.5 ppm) signal on tissue pH. The ghost membranes were prepared according to the method that was modified from previous reports (31,32). Specifically, 15 mL of bovine blood was diluted with an equal volume of 1 × PBS. Then, red cells were separated from plasma by centrifugation (1000 × g, 20 min, 4°C), and the plasma and the buffy coat were removed by aspiration. To remove plasma proteins, the cells were further washed twice with 2 volumes of 172 mM Tris buffer (pH 7.6) by centrifugation (1000 × g, 20 min, 4°C). The washed cells were suspended in isotonic 172 mM Tris buffer (pH 7.6) to an approximate hematocrit of 50% and mixed gently by inversion for approximately 1 min. 10ml aliquots of the above 50% cell suspension were transferred to 50ml centrifuge tubes. 40 ml of 11.35 mM Tris buffer (pH 7.6, 4 °C) was added to the cell suspension and mixed well by stirring for 5 min. Finally, ghost membranes were harvested by centrifugation (20,000 × g, 30 min, 4°C). The ghost membranes were washed ten times with 11.35 mM Tris buffer (pH 7.6, 4 °C) to sufficiently remove proteins and then resuspended in 1× PBS for MRI scans.

Reconstituted phospholipids were used to investigate the dependence of NOE(−3.5 ppm) signal on the chemical composition. Two phantoms containing two types of synthetic phospholipids, DSPC (1,2-Distearoyl-sn-glycero-3-phosphocholine) and DEPC (1,2-Dierucoyl-sn-glycero-3-phosphocholine), were prepared respectively. Note that DEPC (22 × 2 chain length) has a longer aliphatic chain than DSPC (18 × 2 chain length). The reconstituted phospholipid phantoms were prepared by adding the synthetic phospholipids and cholesterol with a molar ratio of 1:1 to the chloroform solution. Chloroform was then removed in the environment of air stream and vacuum. The phospholipids were resuspended in water with the ratio of 1:3 by weight. The samples were then placed in a bath sonicator, and subsequently frozen and thawed for a few cycles to produce liposomes.

### MRI

A continuous wave (CW)-CEST sequence with a 5-second hard saturation pulse followed by single-shot spin-echo echo planar imaging (SE-EPI) acquisitions with a TR of 7 second was used on animal imaging. Images were acquired with matrix size 64 *×* 64, field of view 30 mm *×* 30 mm, 2 mm slice thickness, and one acquisition. A non-imaging CW-CEST sequence with free induction decay (FID) readout and 8-second hard saturation pulse and TR of 10 second was used for phantom imaging. Z-spectra were acquired with RF offsets (Δω) from −10 ppm to 10 ppm. Control images were acquired with RF offsets of 250 ppm. R_1w_ and MT pool size ratio were obtained using a selective inversion recovery quantitative MT method (33). All *in vivo* and phantom imaging were performed at 37°C with saturation power of 0.5 μT. Experiments on all animals and phantoms were performed on a Varian 4.7-T magnet, and the reconstituted phospholipids were performed on a Varian 9.4-T magnet.

### Multiple-pool Lorentzian fit and asymmetric analysis of CEST Z-spectra

We used multiple-pool Lorentzian fitting to process the CEST Z-spectrum from animals, tissue homogenates, and the supernatant. Eq. (1) gives the model function of the Lorentzian fit method.

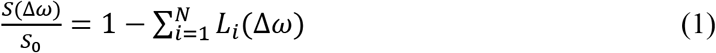

Here, S is the label signal and S_0_ is the control signal; Δ*ω* is the RF frequency offset; *L_i_*(Δ*ω*) = *A_i_*/{1 + (Δ*ω* – Δ_*i*_)^2^/(0.5W_*i*_)^2^}, which represents a Lorentzian line with a central frequency offset relative to the bulk water (Δ_*i*_), peak full width at half maximum (W_*i*_), peak amplitude (A_*i*_); N is the number of fitted pools. A five-pool model Lorentzian fit including the amide, amine, water, NOE(−3.5 ppm), and MT was performed to process the CEST Z-spectra on tissue homogenates and the supernatant. Note that variable frequency offsets of the pools were used in the multiple-pool Lorentzian fit (see Table 1). The NOE(−1.6 ppm) pool was not included in the five-pool model for processing the data from phantoms since the NOE(−1.6 ppm) signals are ignorable in postmortem tissues (34). The fitting was performed to achieve the lowest root mean square (RMS) of residuals between the measured data and the modeled signals. the starting points and boundaries of the amplitude, width, and offset of the fit were list in Table 1. The goodness of the fit was estimated with the sum of RMS of residuals averaged across the whole Z-spectrum.

**Table 1.** Starting points and boundaries of the amplitude (A) and width (W) of all pools in the Lorentzian fit. The unit of peak width is ppm.

|  | Start | Lower | Upper |
| --- | --- | --- | --- |
| $A_{\text{water}}$ | 0.9 | 0.02 | 1 |
| $W_{\text{water}}$ | 1.4 | 0.1 | 10 |
| $A_{\text{amide}}$ | 0.025 | 0 | 0.2 |
| $W_{\text{amide}}$ | 0.5 | 0.4 | 3 |
| $A_{\text{amine}}$ | 0.01 | 0 | 0.2 |
| $W_{\text{amine}}$ | 1.5 | 0.5 | 5 |
| $A_{\text{NOE}(-3.5 \text{ ppm})}$ | 0.02 | 0 | 1 |
| $W_{\text{NOE}(-3.5 \text{ ppm})}$ | 3 | 1 | 5 |
| $A_{\text{semi-solid MT}}$ | 0.1 | 0 | 1 |
| $W_{\text{semi-solid MT}}$ | 25 | 10 | 100 |

The reference signals (S_*ref*_) for the APT, NOE(−3.5 ppm), and semi-solid MT signals were obtained by the sum of all Lorentzians except the corresponding pool estimated (35). An apparent exchange-dependent relaxation (AREX) method (36–38), which inversely subtracts S_lab_ from S_ref_ together with T1 correction, was used to quantify the APT/NOE(−3.5 ppm) effects.

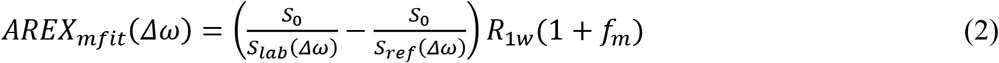

where R_1w_ is the water longitudinal relaxation rate; the subscript ‘mfit’ represents the multiple-pool Lorentzian fit. (1 + *f_m_*) was added to Eq. (2) for all animal imaging to make the inverse subtraction method more specific (38). For all phantom imaging, *f_m_* was set to 0 due to the low MT pool size ratio. The AREX determined semi-solid MT effect was obtained by *R*_1*W*_*L_MT_*/(1 – *L_MT_*), where *L_MT_* is the Lorentzian line for the MT effect (39). The AREXmfit quantified APT and NOE(−3.5 ppm) values were obtained by choosing the maximum value between 4 ppm and 3 ppm in the fitted APT spectra and between −3 ppm and −4 ppm in the fitted NOE(−3.5 ppm) spectra, respectively. The AREX_mfit_ determined MT signal was obtained at −2.3 ppm in the fitted MT spectrum.

We used the asymmetric analysis, in which *S_ref_* was acquired on the symmetrical side relative to the water peak, to process the CEST Z-spectrum from ghost membranes and reconstituted phospholipid samples.

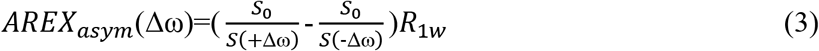

where (+) represents the resonance frequency offset of the labeled protons; (−) is the offset on the opposite side relative to the water peak; the subscript ‘asym’ represents the asymmetric analysis.

### Data analysis and statistics

Regions of interest (ROIs) of tumors, contralateral normal tissues, the corpus callosum and the caudate putamen were delineated in T_2_ weighted images. The corpus callosum and the caudate putamen were selected as representative regions to investigate the NOE(−3.5 ppm) signals in white matter and gray matter, respectively. The student’s t-test was employed to compare the ROI-averaged signals. It was considered to be statistically significant if *P* < 0.05. All data analysis and statistical analyses were performed using MATLAB (Mathworks, Natick, RMA, USA).

## RESULTS

Figure 1 shows the CEST Z-spectra, the corresponding residual spectra as well as the AREXm_fit_ spectra acquired from the homogenates and the supernatant of brain tissues prepared from healthy rats. Figure S1 shows the corresponding magnetization transfer ratio (MTR) spectra in Supplementary information, which is derived from the direct subtraction of S_ref_ and S_lab_ from the multiple-pool Lorentzian fit. The RMS of the residues averaged across the whole Z-spectrum are 0.37% and 0.63% for the homogenates and the supernatant, respectively, which are smaller than the MTR quantified APT signals and NOE(−3.5 ppm) signals from homogenates (APT: 2.16%, NOE(−3.5 ppm): 5.40%) and supernatant (APT: 1.38%, NOE(−3.5 ppm): 0.89%). The low RMS of the residuals indicate high goodness of the multiple-pool Lorentzian fit. Table S1 shows results of the multiple-pool Lorentzian fit from the homogenates and the supernatant in Supplementary information.

**Figure 1.**
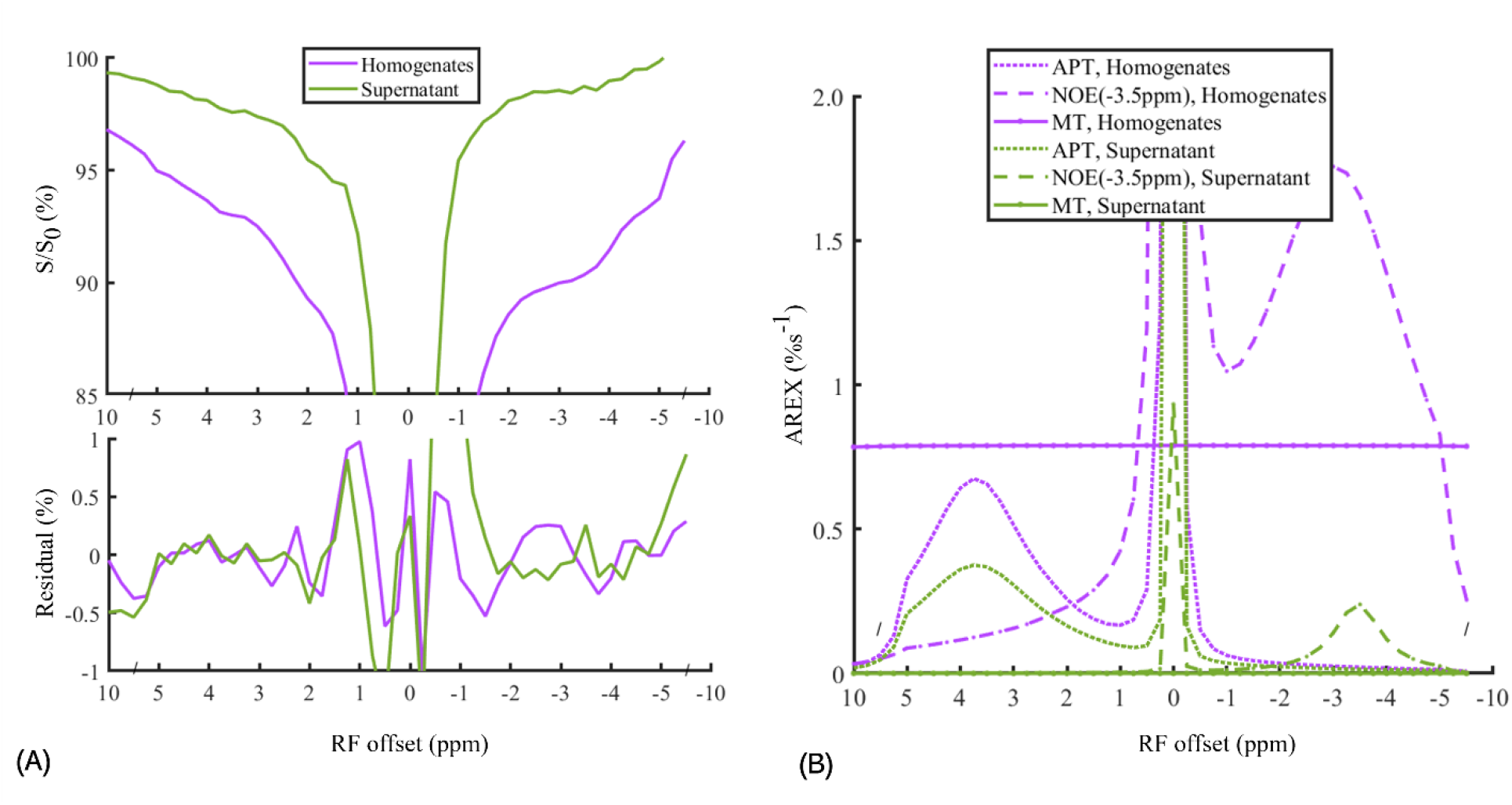
CEST Z-spectra and the corresponding residual spectra (A) and AREX_mfit_ determined APT, NOE(−3.5 ppm), and MT spectra (B) from brain tissue homogenates (purple lines) and the supernatant (green lines). Note that the AREX determined MT in the supernatant is close to 0; the relatively small residuals (< ±0.02%) at ±3.5 ppm compared with the AREX_mift_ determined APT and NOE(−3.5 ppm) signals indicate high goodness of the multiple-pool Lorentzian fitting.

In this study, the ratios of the AREXm_fit_ NOE(−3.5 ppm) signal to the APT signal are derived to assess the relative contributions of proteins and lipids in the brain tissues as below. For the *in vitro* homogenates, the ratio of NOE(−3.5 ppm) signal (1.76 %s^-1^) to APT signal (0.68 %s^-1^) is around 2.59. For the supernatant without lipids, the ratio of the NOE(−3.5 ppm) signal (0.24 %s^-1^) to the APT signal (0.38 %s^-1^) is around 0.63. A reasonable assumption is that the ratio of protein-sourced NOE(−3.5 ppm) signal to protein-sourced APT signal is identical for the homogenates and the supernatant. In this context, the relative contributions of the lipids and proteins to the NOE(−3.5 ppm) signals are derived as around 76% and 24%, respectively. Note that the AREXm_fit_ spectra show no obvious MT effects in the supernatant, which suggests the successful removal of not only semi-solid components but also the membrane lipids, considering that membrane lipids could generate the MT effects (see the Z-spectra beyond ±5 ppm in Figure 2A). The CEST signals at 2 ppm in the tissue homogenates and the supernatant are greater than that measured in the *in vivo* experiments, which may be due to the increased creatine from dephosphorylation of phosphocreatine after the death of the rats (10,40).

**Figure 2.**
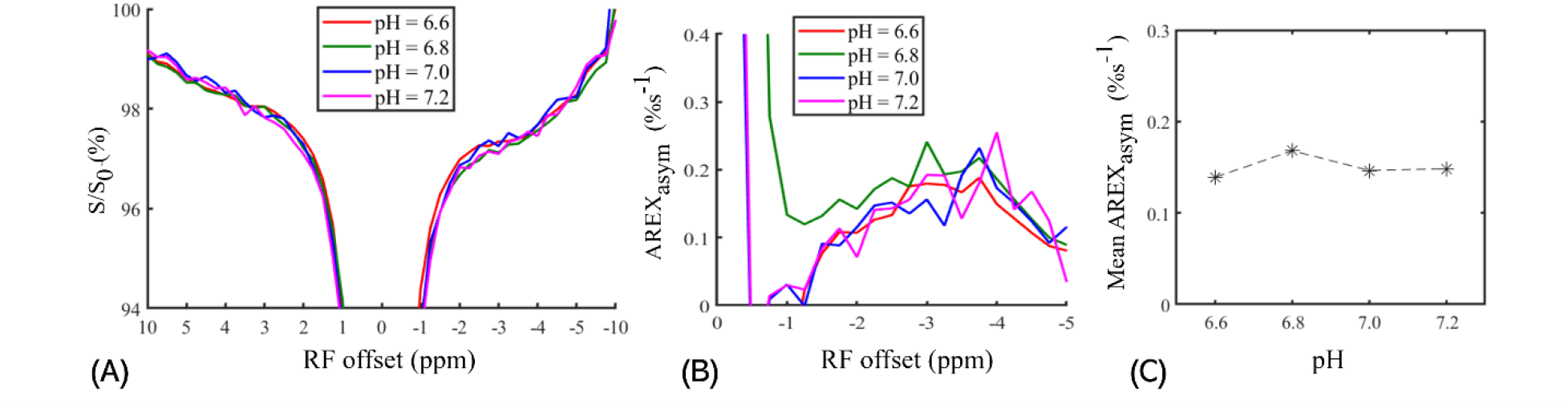
CEST Z-spectra (A), AREX_asym_ spectra (B), and the mean values of AREX_asym_ signals (C) measured from the ghost membrane phantoms with pH of 6.6, 6.8, 7.0, and 7.2. Note that the mean values of AREX_asym_ signals are derived by averaging the AREX_asym_ signals from −5 ppm to −2 ppm in B.

Figure 2 and Figure 3 demonstrate the dependence of the NOE(−3.5 ppm) on two physiological conditions (i.e., pH and the length of aliphatic chain of phospholipid). FigureFigure 2 shows CEST Z-spectra, AREX_asym_ spectra and the mean AREX_asym_ value derived from −5 ppm to −2 ppm measured from the phantom of ghost membranes with varied pH values of 6.6, 6.8, 7.0 and 7.2. The mean AREXasym value derived from −5 ppm to −2 ppm in FigureFigure 2C was used to improve the low SNR of signals measured in the Z-spectra, which demonstrates that the NOE(−3.5 ppm) signal has nearly no dependence on pH. Figure 3 compares CEST Z-spectra and AREX_asym_ spectra measured from DSPC and DEPC. Note that the NOE(−3.5 ppm) signal measured in DEPC is higher than that in DSPC. The aliphatic chain of DEPC is longer than that of DSPC, which may explain its higher NOE(−3.5 ppm) signals.

**Figure 3.**
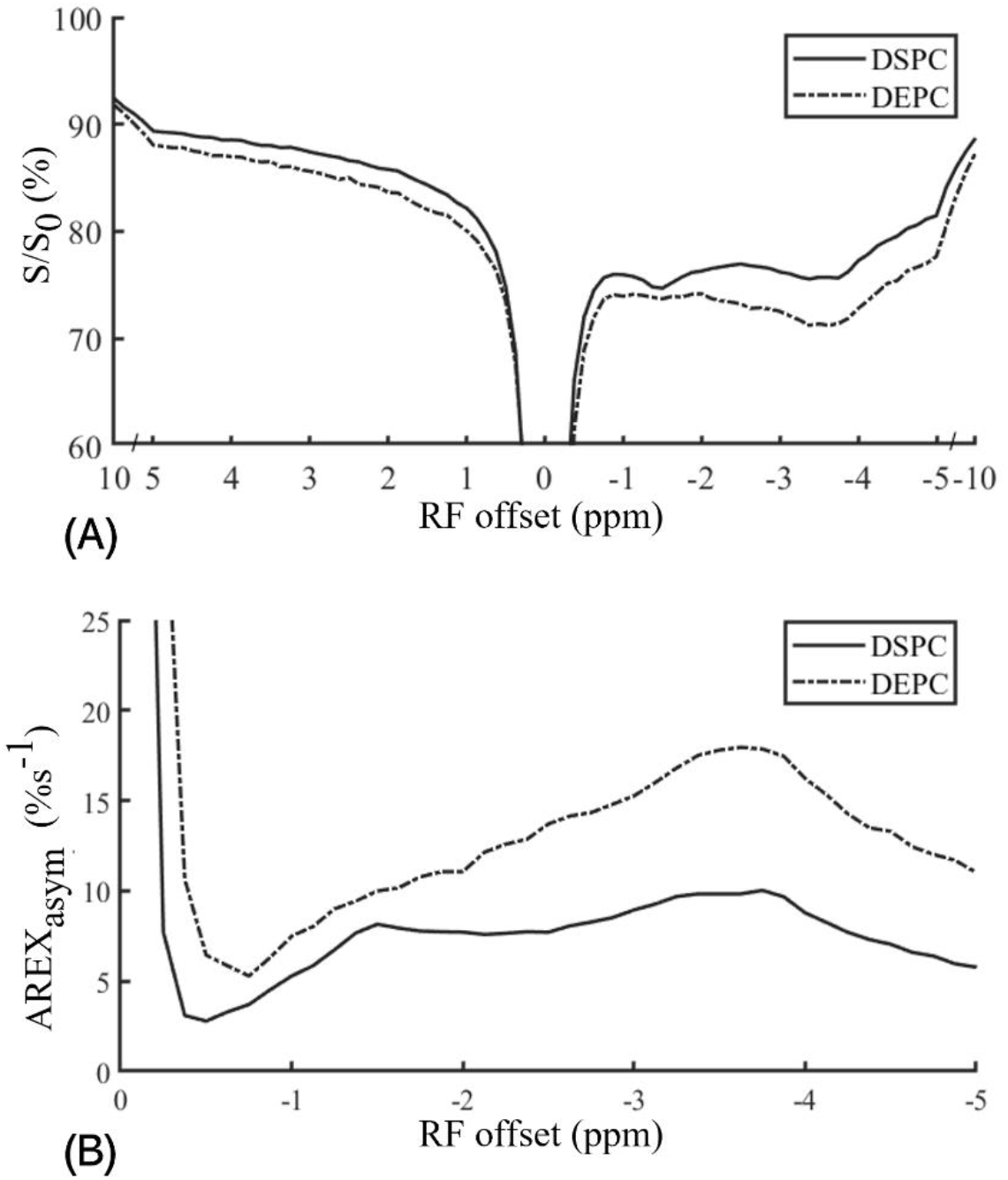
CEST Z-spectra (A) and AREX_asym_ spectra (B) measured from the reconstituted membranes that are prepared with DSPC and DEPC, respectively.

Figure 4 displays the ROI-averaged CEST Z-spectra, the fitting residuals and the AREXmfit spectra acquired from brain tumors and the contralateral normal tissues, as well as from the corpus callosum and the caudate putamen in the brains of rats. The APT signals at 3.5 ppm, the amine signals at 2 ppm, the NOE signals −3.5 ppm and the broad semi-solid MT signals can be clearly observed in the CEST Z-spectra. Figure S2 shows the corresponding MTR spectra in Supplementary information. The RMS of the residues averaged across the whole Z-spectrum are 0.22% and 0.19% for the normal tissues and tumors, respectively, which are much smaller than the MTR quantified APT and NOE(−3.5 ppm) signals from normal tissues (APT: 3.04%, NOE(−3.5 ppm): 8.84%) and tumors (APT: 5.10%, NOE(−3.5 ppm): 8.87%). The RMS of the residues averaged across the whole frequency offset range is 0.18% and 0.21% for the CP and CC, respectively, which are much smaller than the MTR quantified APT signals and NOE(−3.5ppm) signals from CP (APT: 2.82%, NOE(−3.5 ppm): 8.32%) and CC (APT: 2.47%, NOE(−3.5 ppm): 8.47%). The low RMS of the residuals indicate high goodness of the multiple-pool Lorentzian fitting. Table S2 and S3 shows the multiple-pool Lorentzian fitted results from the tumors and contralateral normal tissues as well from the CP and CC in Supplementary information.

**Figure 4.**
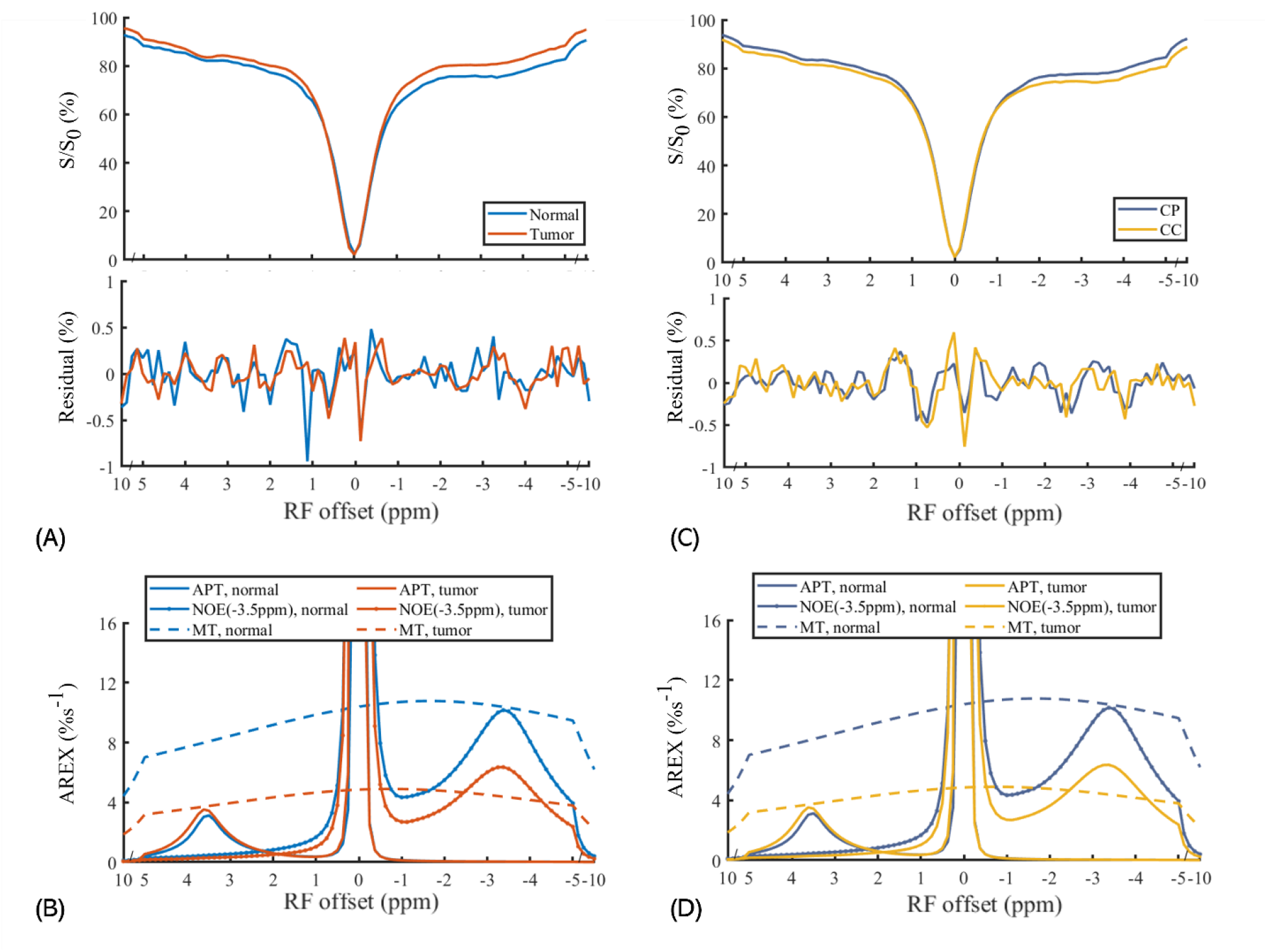
Average CEST Z-spectra and the corresponding residual spectra (A, B) and AREX_mfit_ spectra (C, D) detected from tumors and contralateral normal tissues (left panels), and from the caudate putamen (CP) and the corpus callosum (CC) (right panels). Note that relatively small residuals (< ±0.02%) at ±3.5 ppm compared with the AREX_mift_ determined APT and NOE(−3.5 ppm) signals indicate high goodness of the multiple-pool Lorentzian fitting.

Figure 5 demonstrates maps of T_2_ weighted anatomy, R_1w_, MT pool size ratio, AREXmfit APT at 3.6 ppm, AREX_mfit_ NOE at −3.5 ppm, and AREX_mfit_ MT at −2.3 ppm from a representative rat brain that bears 9L tumors and a representative healthy rat brain. The normal tissues and tumor tissues, the corpus callosum and the caudate putamen in the rat brain are delineated in the T_2_ weighted image for the intuitive comparisons of the image contrasts. Summarily, the maps of T_2_ weighted anatomy and the AREX_mfit_ APT display higher intensities in tumor tissues than in normal tissues while the maps of R_1w_, MT pool size ratio, AREX_mfit_ NOE(−3.5 ppm), and AREX_mfit_ MT show lower intensities; the maps of R1w, MT pool size ratio, AREX_mfit_ NOE(−3.5 ppm), and AREX_mfit_ MT show higher intensities in the corpus callosum than in the caudate putamen. Note that the previous studies have also demonstrated reduced NOE(−3.5 ppm) signals in the cancer tissues (12,17,18,39,41–44), which guarantees the reproducibility of our experiments.

**Figure 5.**
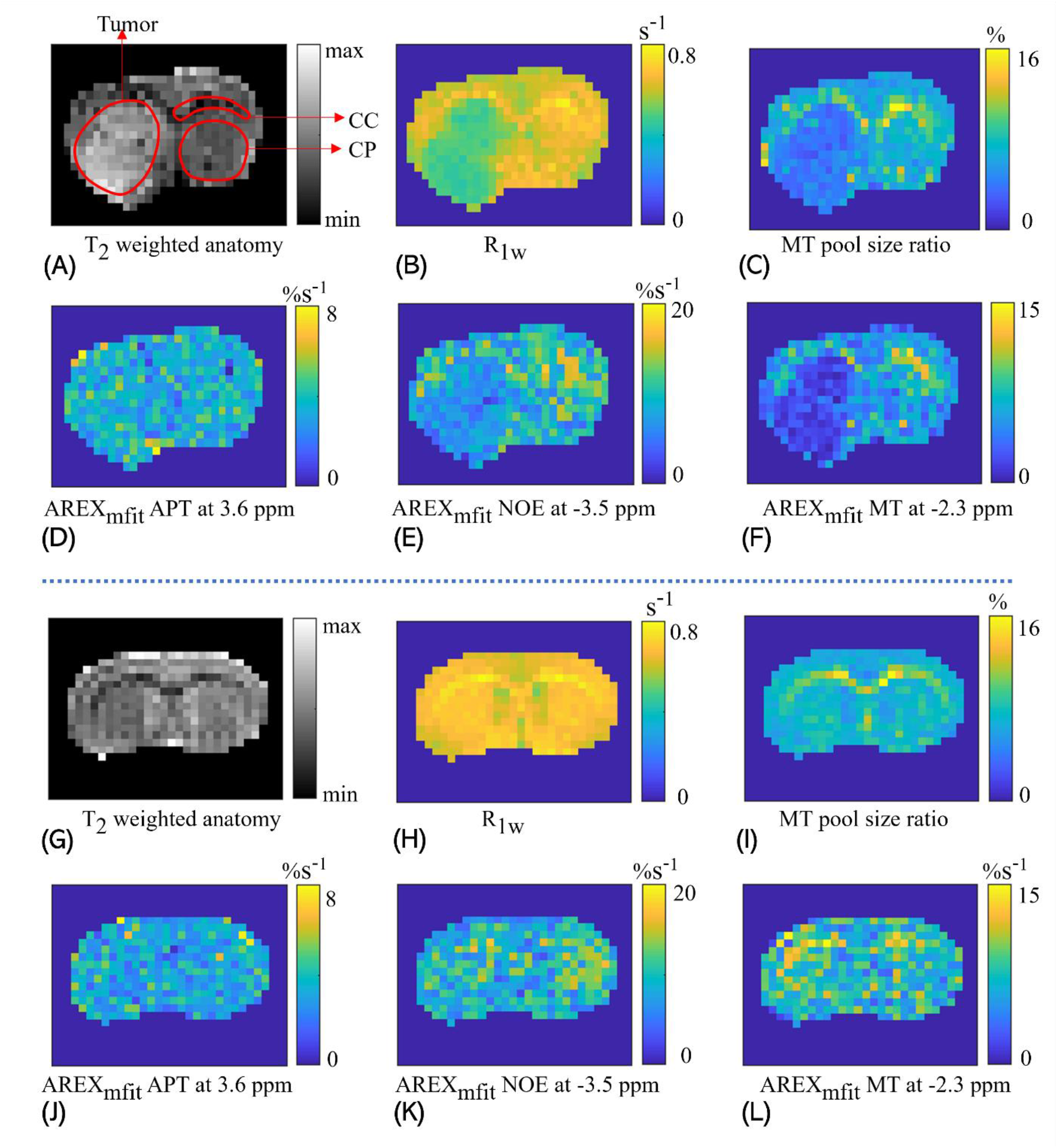
Maps of T_2_ weighted anatomy (A, G), R1w (B, H), MT pool size ratio (C, I), AREX_mfit_ for APT at 3.6ppm (D, J), AREX_mfit_ for NOE(−3.5 ppm) at −3.5ppm (E, K), and AREX_mfit_ for MT at −2.3ppm (F, L), from the brain of a representative rat that bears 9L tumors (A-F) and a representative healthy rat (G-L). In the T_2_ weighted image, the tumor tissues, the caudate putamen (CP) and the corpus callosum (CC) are delineated by red lines.

Figure 6 shows the comparisons of R_1w_, MT pool size ratio, and the quantified CEST signals between tumors and contralateral normal tissues, as well as between the corpus callosum and the caudate putamen. The t-test shows significant differences (*P* < 0.05) of R1w and MT pool size ratio between tumors and normal tissues as well as between the corpus callosum and the caudate putamen; significant differences of the AREX_mfit_ NOE(−3.5 ppm) between tumors and normal tissues, and between the corpus callosum and the caudate putamen; and significant difference of the AREX_mfit_ MT between tumor and normal tissues. To determine the origins of differences in NOE(−3.5 ppm) signals between tumors and normal tissues and between gray matter and white matter, the ratios of the region-averaged AREX_mfit_ signals are derived as below. The ratio of NOE(−3.5 ppm) signal in normal tissues (10.17%) to that in tumors (6.36%) is around 1.60. As inferred above, membrane lipids have a contribution of around 76% to NOE(−3.5 ppm) and proteins have a contribution of around 24%, thus the substantial change in NOE(−3.5 ppm) signal should be mainly explained by lipids. Besides, the ratio of NOE(−3.5 ppm) signal in the corpus callosum (11.85%) to that in the caudate putamen (9.56%) is around 1.24; it also suggestions that the NOE(−3.5 ppm) signals mainly originate from membrane lipids, considering that the white matter (i.e., the corpus callosum) has more membrane lipids and fewer proteins than the gray matter (i.e., the caudate putamen). Note that the AREX_fit_ quantified APT has no significant differences between tumors and normal tissues, which agrees with previous studies (41,43). However, the differences in the MTR quantified APT signals have been observed in other previous studies (45–47), which may be due to confounding factors including R_1w_ and ‘shine through’ effect from direct water saturation and MT that are involved in the MTR metric (48).

**Figure 6.**
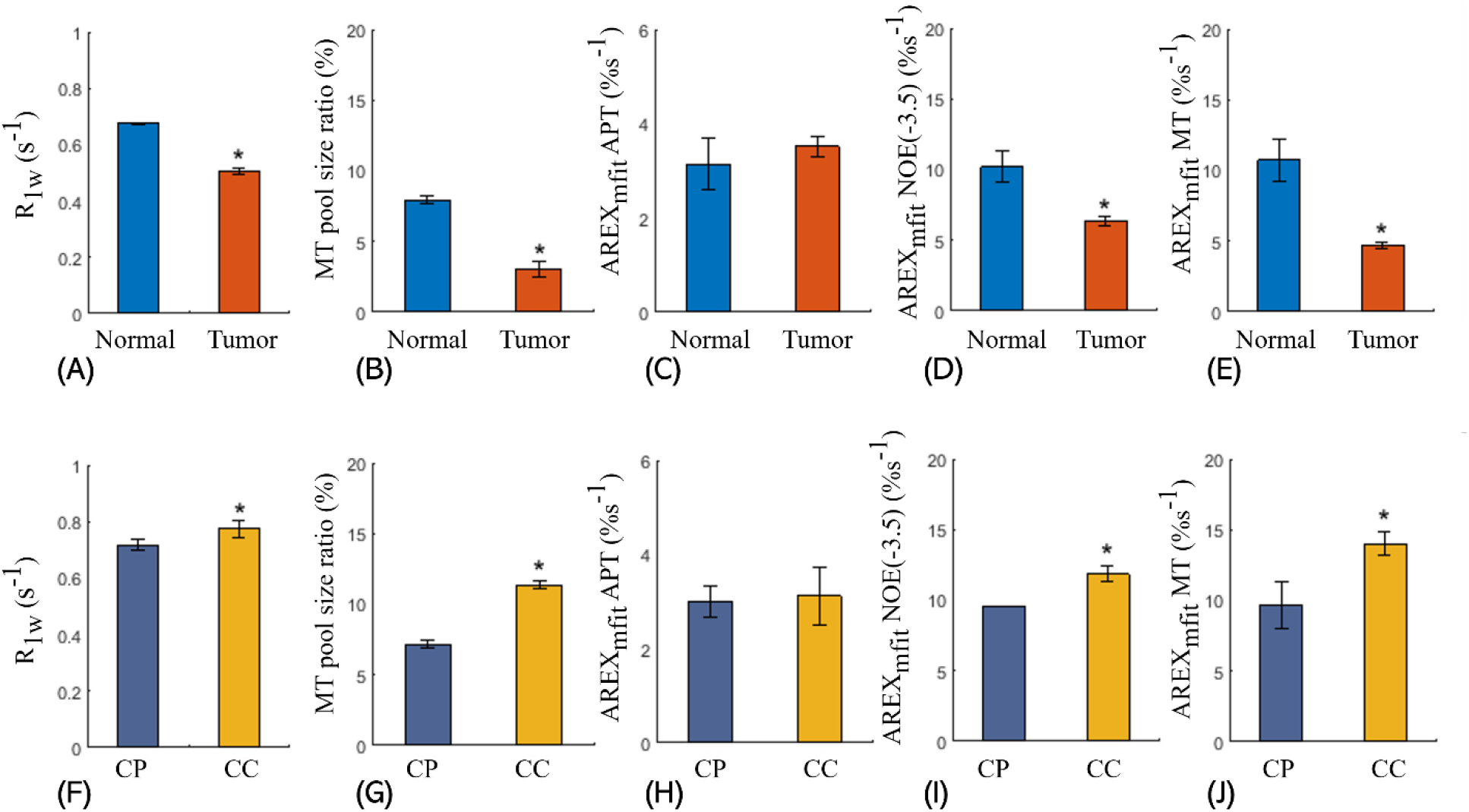
Comparisons between the tumors and the contralateral normal tissues for R_1w_ (A), MT pool size ratio (B), AREX_mfit_ APT at 3.6 ppm (C), AREX_mfit_ NOE(−3.5 ppm) at −3.5 ppm (D), and AREX_mfit_ determined MT at −2.3 ppm (e); and comparisons between and the caudate putamen (CP) and the corpus callosum (CC) for R1w (F), MT pool size ratio (G), AREX_mfit_ determined APT at 3.6 ppm (H), AREX_mfit_ determined NOE(−3.5 ppm) at −3.5 ppm (I), and AREX_mfit_ determined MT at −2.3 ppm (J). The significant differences between two signals are indicated by asterisks (\**P* < 0.05).

## DISCUSSION

In this study, the results of *in vitro* experiments on tissue homogenates and the supernatant of brain tissues suggest that the NOE(−3.5 ppm) signals in the brain mainly originate from membrane lipids. Further studies on ghost membranes and reconstituted phospholipids reveal that the NOE(−3.5 ppm) signals are independent on pH and dependent on the chemical composition of lipids. The independence of lipid-sourced effect on pH along with the similar property of protein-sourced NOE(−3.5 ppm) demonstrated in a recent study (49) can be explained by the mechanism of relayed NOE that NOE is relayed by chemical exchange to achieve saturation transfer from protons on aliphatic chains to the water proton pool (16). Specifically, NOE mediated magnetization transfer is much slower than chemical exchange from relay sites to the water proton pool. Thus, the efficiency of the relayed NOE is dominantly determined by NOE coupling rather than chemical exchange, leading to that the NOE effect is unsusceptible to pH that is associated with the chemical exchange rate. The biophysical mechanism of the dependence on the chemical compositions is hard to be clearly determined in this study, considering that the NOE process is modulated by the chemical composition that varies the number of aliphatic protons and the mobility of molecules in a complex way. The dependence of NOE on the chemical composition is worth a further study since the changes in the chemical composition of membrane lipids are engaged in the pathogenesis of many diseases (50).

The *in vivo* experiments on the rat brain show that the NOE(−3.5 ppm) signals in normal tissues are significantly higher than that in tumors, and the NOE(−3.5 ppm) signals in the corpus callosum (white matter) are significantly higher than that in the caudate putamen (gray matter), which agrees with previous reports (17,20,22,23,42). Given that our *in vitro* experiments demonstrate membrane lipids being the major source of the NOE(−3.5 ppm) signals, the differences in NOE(−3.5 ppm) signals could be largely accounted for by the membrane lipids rather than proteins, which is analyzed as below. The decreases of NOE(−3.5 ppm) signals in tumors should be mainly due to the change in the properties of lipids rather than proteins for the following three reasons. (1) the higher expression of proteins in the tumor tissues (51) could lead to an increase in protein-sourced NOE(−3.5 ppm) signals. (2) since the proteins account for ~24% of NOE signals as inferred in this study, the protein changes in tumors have a limited influence on NOE signals (note that the relative decrease of the NOE signal in tumor is up to 38%); note that a previous study (22,30) reported that the changes in the spatial conformation of proteins could reduce NOE(−3.5 ppm) signals; however, there is no evidence supporting that the spatial conformation of proteins account for the reduced NOE(−3.5 ppm) signals in tumors; (3) numerous reports have demonstrated that membrane lipids in tumors are significantly less rigid than that of normal tissues (51–55); the increased mobility (fluidity) of the cell membranes could reduce the NOE coupling rate, which could explain the reduced NOE(−3.5 ppm) signals in tumors. The higher NOE(−3.5 ppm) signals in white matter than that in gray matter should also mainly result from the variations of membrane lipids, since the dry weight mass fraction of proteins in white matter is 10-15% lower than that in gray matter in proteins while the dry weight mass fraction of lipids in white matter is 60% higher than that in gray matter (56).

The relative contributions of membrane phospholipids and proteins to the NOE(−3.5 ppm) signals are estimated with the reasonable assumption that the ratio of the protein-sourced NOE(−3.5ppm) signal to the protein-sourced APT signal is roughly the same for the homogenates and supernatants. In principle, for the solvent system, the solute (i.e., soluble proteins) is uniformly distributed within the solvent (i.e., water). Hence, the supernatants contain the same soluble proteins as the homogenates, considering that the preparation of supernatants only involves the removal of the insoluble solid components. Note that only soluble proteins could generate the CEST signals with narrow line widths. Also note that there may be some differences in the structure and molecular size of proteins between the supernatant and the homogenates. Considering that the preparation of the supernatant is very quick, the structure and molecular size of proteins should remain unchanged. A previous study indicates that the NOE(−3.5 ppm) signal changes only when there are substantial changes in the protein structure(30). As inferred from the in vitro experiments, membrane lipids have a contribution of around 70% to NOE(−3.5 ppm) and proteins have a contribution of around 30%. In spite of the potential experimental bias mentioned above, a reasonable inference is that the membrane phospholipids have a dominant contribution to NOE(−3.5 ppm) signals.

Besides phospholipids, ghost membranes also contain a large proportion of membrane proteins (57), which have trivial contributions to the APT and NOE(−3.5 ppm) signals. As shown in Figure 2, there are no APT peaks in the Z-spectrum from the ghost membrane phantom, which demonstrates that membrane proteins cannot generate CEST signal peaks with narrow line widths. It may be explained by that the water-insoluble membrane proteins with high molecular weights have limited mobility due to motion restrictions from cell membranes that also have high molecular weights. On the other hand, amide protons inside membrane proteins with large molecular sizes have no access to water protons and thus cannot generate CEST effects(46). Note that phospholipids in cell membranes could generate the NOE(−3.5 ppm) signals with a narrow linewidth since their low molecular weights ensure high mobility.

Protease inhibitor that could inhibit the potential protein degeneration was not used in the preparation of the homogeneities in this study since the most widely used inhibitor (e.g., ethylenediaminetetraacetic acid) has a cation (Ca++/Mg++ chelator) that could change the structures of proteins (i.e., degeneration of proteins) and thus alter the NOE signals as well as the CEST signals. To avoid the degradation of proteins due to proteases, the preparation of tissue samples was performed within a short time at a low temperature (4°C) to minimize protease activities. In addition, MRI measurements of these samples were also performed within 0.5 h to reduce the protease-induced decomposition of proteins.

The multiple-pool Lorentzian fit was used to quantify the CEST, NOE and MT signals, in which the limitations of this method could introduce the relevant bias in the study. However, the relevant bias of the fitting can be reduced to a low level under the following specific conditions. First, although the semi-solid MT can be better fitted by super-Lorentzian absorption line shape (58–60), we fit CEST signals obtained with RF offsets within ±10 ppm, where the semi-solid MT can be fitted by the Lorentzian function with small errors (61). In biological tissues, besides the five pools used in the multiple-pool Lorentzian fit, the amines at 3 ppm and the hydroxyls from 1 ppm to 3 ppm also contribute signals in the Z-spectrum. However, by observing the CEST Z-spectrum, these pools can be roughly treated as a single pool centered at 2 ppm. Previous studies have shown that the APT can be still quantified at relatively low saturation power with the presence of fast exchanging amines at 3 ppm (41). In addition, aliphatic NOE signals include many peaks spanning from −5 ppm to −1 ppm. However, by observing the CEST Z-spectrum, these pools can be roughly treated as two peaks at around −1.6 ppm and −3.5 ppm (12). The CEST signals at 3.5 ppm may consist of the APT and the aromatic NOE (62), both of which should originate from proteins.

In the brain tissues, lipids can be cataloged into three types: myelin lipids, membrane phospholipids, and neutral lipids (56). Myelin lipids, that are the major components of the myelin sheath in nerve fibers in white matter, have the lowest mobility of the three types of lipids, which can be manifested by its very broad line width observed in the conventional MT Z-spectrum. Neutral lipids have a very low concentration in brain (50), which exist in the form of lipid droplets as energy reserves in bio tissues and cannot produce CEST effects. Note that neutral lipids in tissues could generate artifacts in Z-spectrum, which need to be carefully removed in CEST imaging (63–65). The NOE(−3.5ppm) signals arise from the aliphatic chains of phospholipids. Note that the NOE(−1.6 ppm) is another NOE effect, which is sourced from the choline head group of phospholipids (12,34,42,66,67). To date, myelin lipids can be detected by MT imaging (14,15), and neutral lipids can be detected by magnetic resonance imaging with a postprocess of fat-water separation (68), and magnetic resonance spectroscopy (MRS) (69). However, the *in vivo* detection of membrane lipids is challenging because of the short transverse relaxation time of the membrane lipids. In this study, we demonstrate that NOE(−3.5 ppm) imaging could serve as an MRI tool for monitoring membrane phospholipid.

In this preliminary work, although we investigated the partial properties of the membrane-lipid-sourced NOE(−3.5 ppm) signals (i.e., regarding the chemical composition and the pH), its complete biophysical properties warrant further systematic studies that are could be beneficial to interpreting its underlying contrast mechanism of membrane lipids associated with the pathogenesis of diseases at molecular levels. The length of aliphatic chains of lipid molecule could change the number of protons and thus affect the NOE signals via a concentration effect. Besides, it also alters the fluidity of membrane (70) and thus modulate the efficiency of the NOE. In addition, the unsaturation of membrane lipids, the concentration of cholesterol in the membrane and the behavior of the membrane-bound proteins could also modulate the efficiency of the NOE by altering the fluidity of membrane (71–73). Note that the changes in the chemical composition of membrane lipids in the pathogenesis of diseases are companied with multiple factors mentioned above (50,74). In future studies, univariate experiments based on reconstituted membrane phantoms can be used to investigate these factors, separately.

## CONCLUSION

In this study, we demonstrate that the NOE(−3.5 ppm) signals in the brain mainly originate from membrane lipids, which suggests that membrane lipids account for the origins of the NOE(−3.5 ppm) contrast differences between tumors and normal tissues, and between gray matter and white matter. NOE(−3.5 ppm) imaging could be exploited as a new MRI method to monitor membrane lipids *in vivo*, which has great potential for clinical applications.

